# *C. elegans* REMO-1, a glial GPCR, regulates stress-induced nervous system remodeling and behavior

**DOI:** 10.1101/2020.06.03.127894

**Authors:** In Hae Lee, Carl Procko, Yun Lu, Shai Shaham

## Abstract

Animal nervous systems remodel following stress. Although global stress-dependent changes are well documented, contributions of individual neuron remodeling events to animal behavior modification are challenging to study. In response to environmental insults, *C. elegans* become stress-resistant dauers. Dauer entry induces amphid sensory-organ remodeling, in which bilateral AMsh glial cells expand and fuse, allowing embedded AWC chemosensory neurons to extend sensory receptive endings. We show that amphid remodeling accelerates dauer exit upon exposure to favorable conditions, and identify a G protein-coupled receptor, REMO-1, driving AMsh glia fusion, AWC neuron remodeling, and dauer exit. REMO-1 is expressed in and localizes to AMsh glia tips, is dispensable for other remodeling events, and promotes stress-induced expression of the remodeling receptor tyrosine kinase VER-1. Our results demonstrate how single-neuron structural changes affect animal behavior, identify key glial roles in stress-induced nervous system shape changes, and demonstrate that remodeling primes animals to respond to favorable conditions.

## INTRODUCTION

Nervous systems are robust in connectivity (Goodman and Shatz, 1993), allowing for reproducibility in behavior, yet malleable in function, permitting adaptation to fluctuating environments (Albert and Riddle, 1983; Bourne and Harris, 2008; Cuschieri and Bannister, 1975; Engström, 1967; Flock et al., 1999; Golden and Riddle, 1984; Goodman and Shatz, 1993; Murray, 1993; Procko et al., 2011). Such functional plasticity can be accompanied by structural alterations. For example, in response to dehydration or lactation, astrocytes retract their synapse-associated processes in the paraventricular and supraoptic nuclei to alter synaptic activity (Chapman et al., 1986; Gregory et al., 1980; Hatton et al., 1984; Theodosis et al., 1986; Theodosis and Poulain, 1984, 1989). Similarly, environmental stimuli can remodel sensory organs. For example, in the organ of Corti of the mammalian inner ear, glia-like Deiters’ cells form a scaffold that supports outer hair cells (Rio et al., 2002). Upon exposure to high intensity sound, Deiters’ glia are displaced towards the outer hair cells to reduce cochlear sensitivity, protecting against hearing loss (Flock et al., 1999).

Changes in animal behavior have been correlated with large-scale changes in glia and neuron ensembles, and plasticity mechanisms in individual neurons have been extensively explored (Citri and Malenka, 2008). However, directly linking changes at single synapses or at single sensory receptive endings to animal behavior, a major goal for understanding neural processing, is challenging in animals where many neurons contribute to behavior output.

The nematode *C. elegans* is an excellent model in which to study sensory organ remodeling and single-cell contributions to animal behavior. Upon exposure to stressful conditions, including high temperature, starvation, and crowding, animals enter an alternative developmental state called dauer (Cassada and Russell, 1975; Golden and Riddle, 1984), in which morphological alterations in many organs are observed. The sensory nervous system of dauer animals undergoes striking changes in morphology and gene expression. These changes have been primarily studied in the context of the amphid and inner labial sensilla (Albert and Riddle, 1983; Peckol et al., 2001; Schroeder et al., 2013). Amphids are the primary sensory organs of *C. elegans*, and detect chemosensory, osmotic, mechanical, thermal, light, and other stimuli (Bargmann and Mori, 1997; Perkins et al., 1986; Troemel, 1999; Troemel et al., 1995). Each of the bilateral amphids consists of 12 sensory neurons and two glial cells, the AMsh and AMso glial cells, which wrap around sensory neuron receptive endings (NREs) (Ward et al., 1975). Under normal growth conditions, the NRE of each of the bilateral AWC amphid neurons, which houses receptors for volatile odorants (Troemel, 1999; Troemel et al., 1995), is individually ensheathed by processes of adjacent AMsh glial cells (Figures 1A and 1B). Upon dauer entry, AWC NREs expand in lock-step with their associated AMsh glia wrappings. In half of dauer animals, the bilateral AMsh glia fuse, allowing AWC NREs to expand beyond the midline (Figure 1C) (Albert and Riddle, 1983; Golden and Riddle, 1984; Procko et al., 2011). Thus, AMsh glia delimit AWC NRE growth. Ablation studies demonstrate that AMsh glia remodel independently of AWC NREs (Procko et al., 2011). The functional consequences of AMsh glia and AWC neuron alterations in dauers are not known.

**Figure 1.**
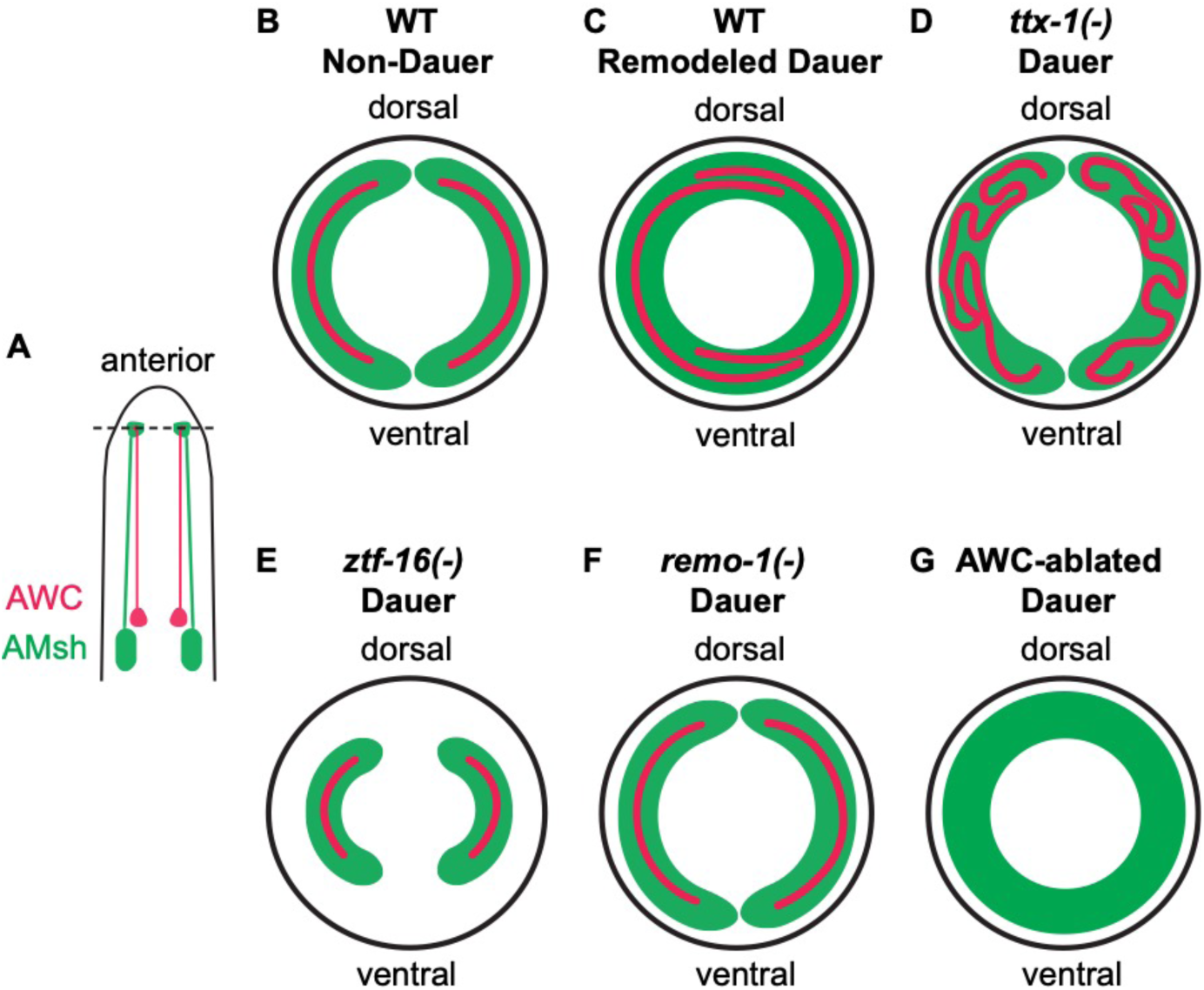
Schematic of amphid sensory organ remodeling in *C. elegans* dauer animals. (A) Schematic of the head of the animal, showing the two bilateral AMsh glia (green) and AWC sensory neurons (red). The horizontal dashed line indicates the position of the transverse sections shown in B-G. (B,C) Sections through the nose tip showing the relative positions of the AWC neuron receptive ending and the ensheathing AMsh glia in wild-type non-dauer (B) and remodeled dauer (C) animals. Note AMsh glial fusion and AWC receptive ending expansion and overlap in dauers. (D-G) Sections through the nose tip showing the relative positions of the AWC neuron receptive endings and the ensheathing AMsh glia in *ttx-1(p767)* (D), *ztf-16(ns171)* (E), *remo-1(ns250, −820, −821)* (F), and AWC-ablated (G) dauer animals. Not to scale. (D,E) Based on (Procko et al., 2011).

Previous studies from our lab identified several AMsh glia proteins implicated in remodeling. These include the cell-fusion protein AFF-1; a VEGFR-related tyrosine kinase protein, VER-1; the Otd/Otx transcription factor TTX-1; and the zinc-finger transcription factor ZTF-16 (Procko et al., 2011; Procko et al., 2012). *ver-1* expression in AMsh glia is induced by cultivation at high temperature (25°C) or by dauer entry, and requires direct binding of TTX-1 to *ver-1* regulatory sequences (Procko et al., 2011).

Here we use a novel assay to demonstrate that amphid remodeling is required for timely dauer exit following exposure to a favorable environment. We identify a gene, which we have named *remo-1* (*<u>remo</u>deling defective-1*), and which encodes a putative 7-transmembrane G protein-coupled receptor (GPCR) that is required for dauer-induced AMsh glia and AWC neuron remodeling. REMO-1 protein is required for expression of VER-1 in AMsh glia, functions in and is expressed in AMsh glia, and localizes to the tip of the amphid sensilla. Importantly, loss of *remo-1* delays dauer exit. Our studies directly link shape changes in a single neuron-glia pair to animal behavior, and demonstrate that glia-dependent nervous-system remodeling following stress allows animals to efficiently detect restoration of a non-stressful environment.

## RESULTS

### Glial REMO-1 regulates stress-induced VER-1/RTK expression

To uncover genes required for AMsh glia and AWC neuron remodeling in dauer animals, we sought mutants that fail to induce *ver-1* promoter::GFP transgene expression in adults when cultivated at 25°C (Procko et al., 2011). Since *ver-1* is also up-regulated in dauer animals, and is implicated in AMsh glia remodeling (Procko et al., 2011), we reasoned that our screen might identify remodeling genes. Indeed, roles for *ttx-1* and *ztf-16* were discovered in this way (Procko et al., 2011; Procko et al., 2012). From this screen, we identified a mutant, with designated allele number *ns250*, that fails to turn on P*ver-1*::GFP in young adults cultivated at 25°C and in dauer larvae induced by starvation (Figures 2A and 2B). Genetic mapping and whole-genome sequencing studies show that *ns250* is not an allele of previously-identified remodeling genes. Rather, *ns250* mutants carry a lesion in the gene *F47D2*.*11*, which we have renamed *remo-1* (*<u>remo</u>deling defective<u>-1</u>*). *remo-1* is a putative Srz*-*type G protein-coupled receptor (GPCR; Figure 2C), and is conserved in other nematode species (Figure S1) (Robertson and Thomas, 2006). The functions of nematode-specific Srz GPCRs are not well understood. The *remo-1(ns250)* allele is predicted to cause a glutamic acid to lysine change at position 278 (E278K) in the C-terminal intracellular tail of the protein (Figure 2C). This residue is part of a highly conserved threonine-glutamic acid-isoleucine (TEI) motif conserved in a subfamily of Srz GPCRs (Figure S1), and lies at the terminal membrane-cytosol interface (Figures 2C and S1), suggesting that its mutation could lead to deficits in REMO-1 function.

**Figure 2.**
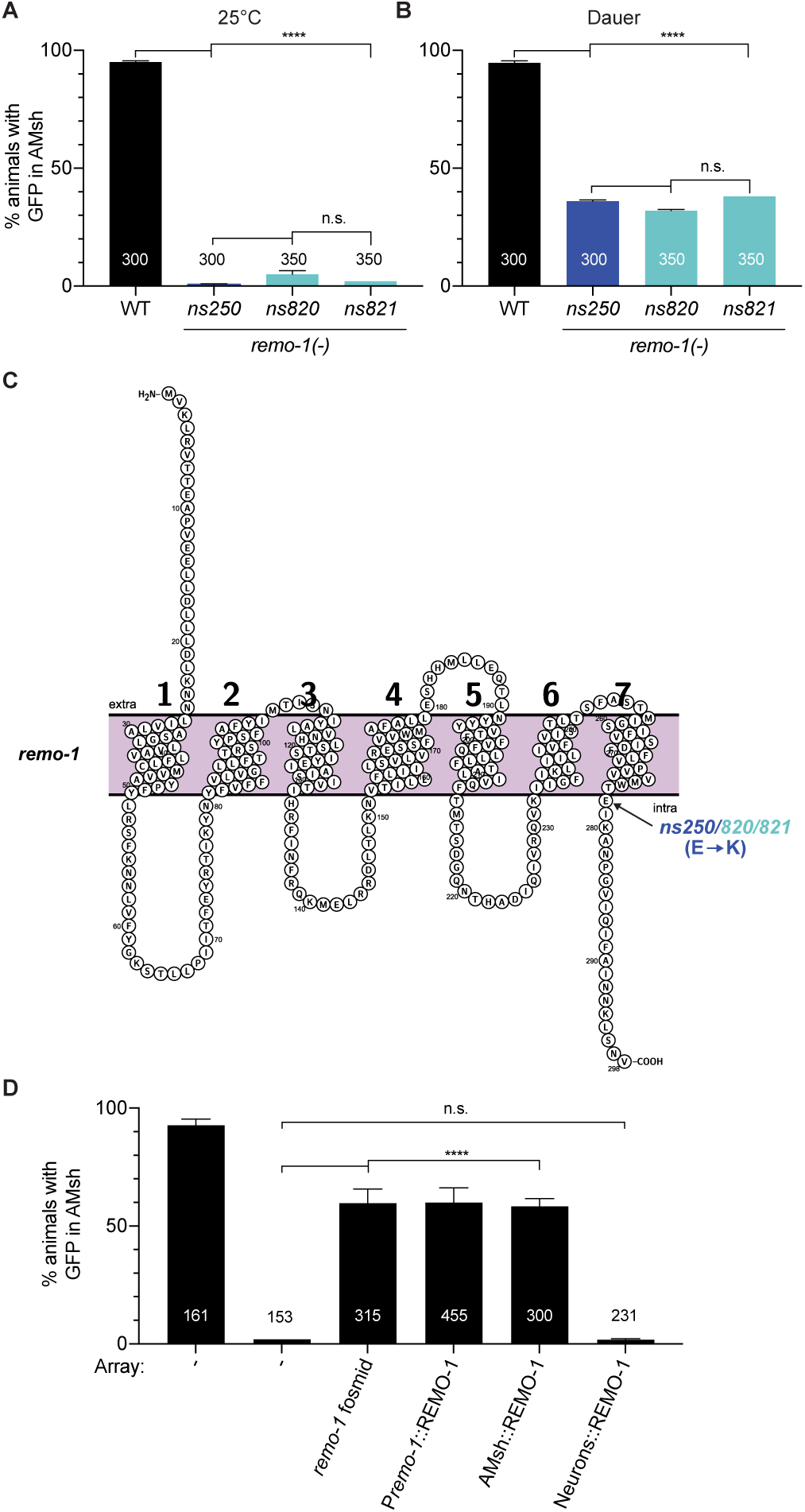
Glial REMO-1 regulates stress-induced VER-1/RTK expression. (A,B) P*ver-1*::GFP expression in the AMsh glia in indicated genotypes at 25°C in young adult (A) or in dauer larvae induced by starvation (B). WT, wild-type. The numbers of animals scored is inside bars. Error bars, SEM of three independent scoring experiments. n.s., p > 0.05, ****, p < 0.0001. (C) Diagram of REMO-1 predicted protein structure and mutation sites generated using Protter software. *ns250/820/821*: glutamic acid to lysine change at position 278 (E278K) indicated by arrow; (D) Histogram detail as in (A,B) All strains are *remo-1(ns250)* mutant background, except for wild-type animals on the left. Rescue experiments conducted with genomic fosmid rescue construct WRM0638cF08, which spans the *remo-1* locus; a smaller *remo-1* genomic fragment; an AMsh glia promoter (*F16F9*.*3*) driving *remo-1* cDNA; or amphid sensory neuron promoter (*dyf-7*) driving *remo-1* cDNA. All rescue constructs were injected with a co-injection marker (P*unc-122*::RFP). Animals screened at young adult stage. Three independent lines were observed for each rescue construct.

To confirm that the E278K change is indeed the causal mutation in *remo-1(ns250)* mutants, we utilized CRISPR to recreate the same genomic lesion in otherwise wild-type animals carrying the *ver-1*::GFP transgene. Two independent mutants, *remo-1(ns820)* and *remo-1(ns821)*, were isolated, and both fail to induce P*ver-1*::GFP expression in young adults cultivated at 25°C and in dauer larvae induced by starvation (Figures 2A and 2B). Furthermore, transgenes containing a genomic fosmid clone encompassing the *remo-1* locus, or a smaller genomic fragment, restore P*ver-1*::GFP expression at 25°C to *remo-1(ns250)* mutants (Figure 2D). Finally, we found that expression of *remo-1* cDNA using an AMsh glia-specific promoter, derived from the *F16F9*.*3* gene, but not an amphid sensory-neuron promoter, derived from the *dyf-7* gene, also restores P*ver-1*::GFP induction to *remo-1(ns250)* mutants (Figure 2D). Taken together, these results strongly support a role for *remo-1*, acting cell-autonomously in AMsh glia, in the induction of *ver-1* expression.

### REMO-1 functions in AMsh glia to regulate dauer-induced AMsh glia remodeling

We next sought to determine whether *remo-1* is required for remodeling of AMsh glia in dauer animals. Remodeling is difficult to follow using conventional or even super-resolution fluorescence microscopy, as the structures in question are often separated by <20 nm. We therefore functionally monitored remodeling by assessing the frequency of fusion of the two AMsh glia, using a validated cytoplasmic mixing assay we previously developed (Figure 3A) (Procko et al., 2011). Briefly, *daf-7(e1372)* single-or *daf-7(e1372)*; *remo-1(ns250)* double-mutants, which, independently of crowding or starvation, become dauers at 25°C as a result of the *daf-7(e1372)* lesion, were engineered to carry an AMsh glia::GFP transgene from an unstable extra-chromosomal array (*nsEx1391*). First-stage larvae (L1) of these strains that express GFP in only one of the bilateral AMsh glial cells were then isolated. These mosaic animals were cultivated at 25°C for at least 48 hours to induce dauer entry, and scored for redistribution of GFP to both cells, indicating fusion. While 46% of *daf-7(e1372)* single mutants redistribute GFP to both AMsh glia, consistent with our previous studies (Procko et al., 2011; Procko et al., 2012), no redistribution is seen in *daf-7(e1372)*; *remo-1*(*ns250*) double mutants (Figure 3B). Similarly, the CRISPR-generated strains, *remo-1(ns820)* and *remo-1(ns821)*, also show no GFP redistribution (Figure 3B). Consistent with these findings, *daf-7(e1372)* dauer animals that fail to redistribute GFP, do not show fusion when examined by electron microscopy (EM) serial reconstructions (n=3), while animals that redistribute GFP, do (n=2; Figures 3C and 3D); and EM reconstructions of *daf-7(e1372)*; *remo-1(ns250)* double mutants reveal lack of fusion (n=2; Figure 3E). Importantly, as with P*ver-1*::GFP rescue, expressing REMO-1 in AMsh glia, but not in AWC and other amphid neurons, is sufficient to fully restore AMsh glia fusion to *remo-1(ns250)* mutants (Figure 3B). Thus, REMO-1 is required in AMsh glia for dauer-dependent AMsh glia remodeling, and functions downstream of or in parallel to DAF-7/TGFβ.

**Figure 3.**
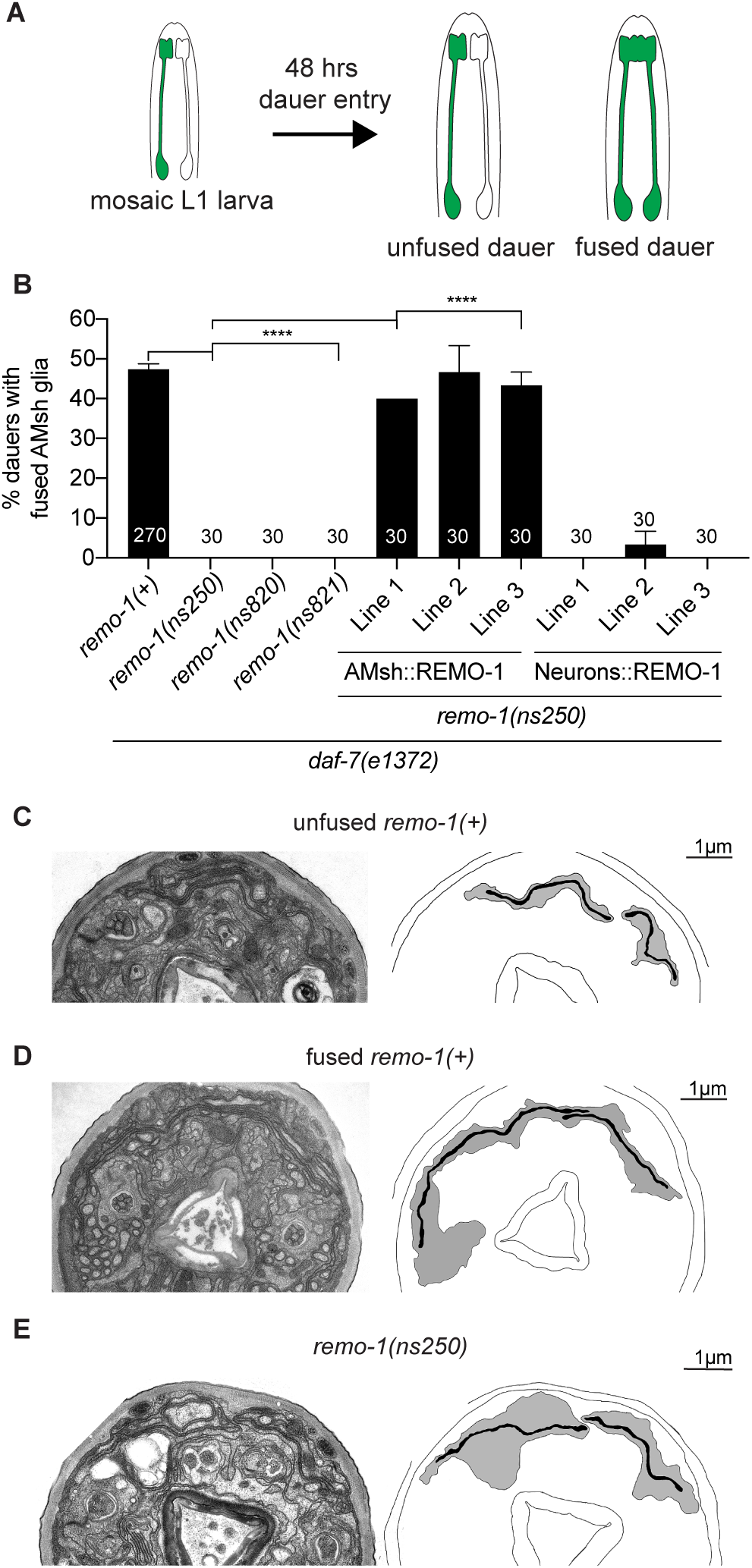
REMO-1 functions in AMsh glia to regulate dauer-induced AMsh glia remodeling. (A) Cytoplasmic mixing assay. Schematic of the head of the animal which expresses an AMsh::GFP transgene from an unstable extrachromosomal array (*nsEx1391*). First-stage mosaic larvae expressing GFP in one of the two AMsh glia were selected and cultivated for at least 48 hours at 25°C. If the two AMsh glia do not fuse, the mosaic animals continue to express GFP in only one of the two glia. If the two AMsh glia undergo fusion, cytoplasmic mixing occurs and both cells fluoresce. Entry into dauer is facilitated by *daf-7(e1372)* temperature-sensitive allele. Anterior is up. Adapted from (Procko et al., 2011). (B) Percentage of animals with fused AMsh glia as scored by cytoplasmic mixing assay in indicated genotypes. All strains carry the *daf-7(e1372*) mutation to induce dauer when cultivated at 25°C. Cell-specific expression of *remo-1* in AMsh glia (*F16F9*.*3* promoter) rescues dauer-induced AMsh glia fusion, whereas expression in the amphid neurons (*dyf-7* promoter) does not. Rescue experiments were conducted in *remo-1(ns250)* mutant background and three independent extrachromosomal arrays were observed per rescue construct. Number of animals scored is inside bars. Error bars, SEM of at least three independent scoring experiments. ****, p < 0.0001. (C-E) Representative electron micrographs (EM) and schematic outlines of amphid sensory organs in dauer animals of given genotypes. Dauer larvae were induced by the *daf-7(e1372)* mutation cultivated at 25°C. Scale bar, 1μm.

### REMO-1 is expressed in AMsh glia and localizes to the glial tip

The P*ver-1*::GFP and AMsh glia fusion rescue studies suggest that REMO-1 may be expressed in AMsh glia. To assess this, we generated wild-type animals carrying a bicistronic transgene consisting of the entire *remo-1* genomic locus, including its 5’ intergenic region, and DNA encoding mKate2 fluorescent protein tagged with two nuclear localization signals, separated by a trans-splicing site (Figure 4A). We found that mKate2 expression is detected in AMsh glia in all larval stages, as confirmed by co-expression with a pan-glial P*mir-228*::myristoylGFP transgene (Figure 4B). We also detected rare expression in the ASI and ASJ neurons (Figure S2), confirmed by DiO staining, but never in AWC neurons. While ASI and ASJ control whole-animal dauer entry and exit (Bargmann and Horvitz, 1991; Cornils et al., 2011), *remo-1* mutants can form dauers at normal rates, and *remo-1* AMsh and AWC remodeling defects are not rescued by expression of *remo-1* in these neurons (Figure 3B).

**Figure 4.**
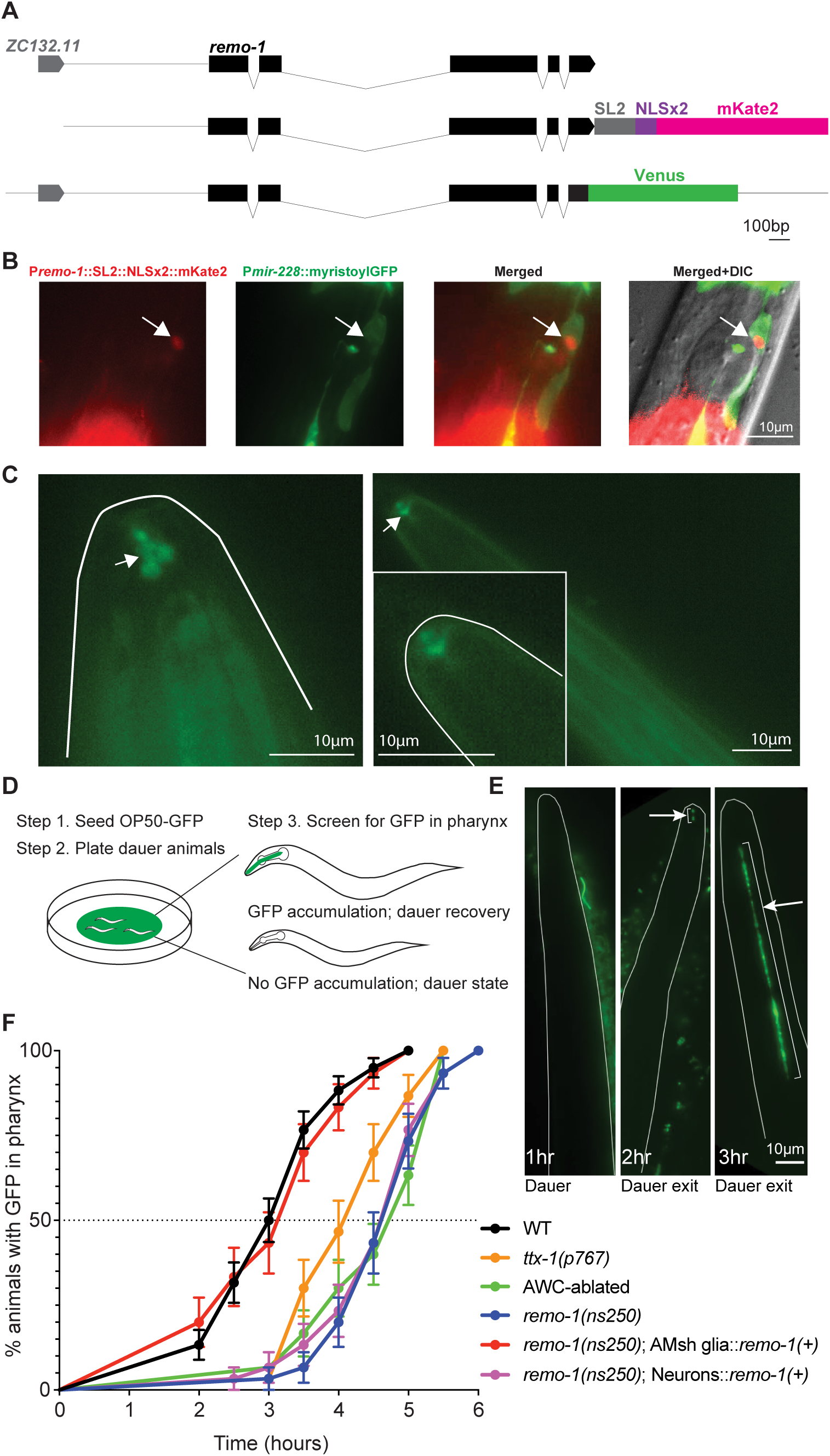
REMO-1 is expressed and localized at the tip of the AMsh glia to regulate dauer exit timeline. (A) Top: schematic of *remo-1* and its upstream gene, *ZC132*.*11*. Middle: *remo-1* transcriptional reporter containing a bicistronic transgene consisting of the entire *remo-1* genomic locus, including its 5’ intergenic region, followed by mKate2 fluorescent protein tagged with two nuclear localization signals, separated by a trans-splicing site. Bottom: *remo-1* translational reporter using CRISPR to knock-in a transgene encoding Venus fluorescent protein just upstream of the *remo-1* stop codon. Black boxes represent exons, black lines indicate introns, grey line indicates 5’ intergenic region. Scale bar length 100 bp. (B) Fluorescence images depicting expression of *remo-1* transcriptional reporter co-expressed with a pan-glial P*mir-228*::myristoylGFP transgene. *remo-1* promoter expression in AMsh glia (left, arrow); *mir-228* pan-glial promoter expression (middle left, AMsh glia arrow); merged (middle right; AMsh glia arrow); merged with DIC (right, AMsh glia arrow). (C) Fluorescence images depicting subcellular localization of REMO-1 in AMsh glia (arrow) in adult (left) and dauer (right) animals. REMO-1 accumulates at the tip of the amphid region. Adult animals were cultivated at 25°C and dauer state was induced by starvation. Magnified inset of the Venus accumulation at the tip of the nose. (B,C) Scale bar, 10μm. (D) Dauer recovery assay. Dauer animals induced by starvation were plated on agar plates with GFP-tagged OP50 bacteria lawn at 15°C. Dauer animals were screened for dauer exit by the onset of GFP accumulation in the pharynx. (E) Fluorescence images showing timeline of GFP-tagged OP50 bacteria accumulation in the pharynx of dauer animals cultivated at 15°C. Images are maximum z-stack projections of the entire head volume of the animal. Outline of the animal is marked in grey. Arrow marks the accumulation of GFP-tagged OP50 bacteria in the pharynx. Anterior, up. Time in hours following exposure to GFP-tagged OP50 bacteria. Each image is of a different animal. Scale bar, 10μm. (F) The rates of dauer exit measured by the onset of GFP-tagged OP50 bacteria accumulation in the pharynx. Black dotted line indicates the time-point in which 50% of the given population recovers pumping. *ttx-1(p767)*, AWC-ablated, and *remo-1(ns250)* mutants show significant dauer exit delay compared to wild-type (p<0.0001). Dauer exit delay in *remo-1(ns250)* is rescued by restoring WT REMO-1 in AMsh glia (*F16F9*.*3* promoter) but not in amphid sensory neurons (*dyf-7* promoter). Dauer larvae induced by starvation.

To examine REMO-1 subcellular localization, we used CRISPR to knock in a transgene encoding Venus fluorescent protein sequences just upstream of the *remo-1* stop codon (Figure 4A) (Arribere et al., 2014). Transgenic animals show consistent Venus expression in two adjacent regions at the tip of the nose in both dauer and non-dauer larvae (Figure 4C), consistent with localization at the site of AMsh glia and AWC remodeling events.

### REMO-1 is also required for AWC neuron remodeling

Since AMsh glia can affect sensory NRE morphology (Procko et al., 2011; Procko et al., 2012; Singhvi et al., 2016), we assessed whether REMO-1 regulates AWC remodeling using EM. In *ttx-1* mutant dauers, AWC neurons can expand within the limited space provided by each unfused AMsh glial cell by forming membrane loops to accommodate excess AWC growth (Figure 1D) (Procko et al., 2011). In these animals, AWC and AMsh glia remodeling are, therefore, uncoupled. In *ztf-16* mutants, AMsh glia and AWC NREs are not well developed in non-dauer animals and remain the same in dauer animals (Figure 1E) (Procko et al., 2012). In *remo-1(ns250, ns820*, or *ns821)* mutants, we found that AWC dendrites fill the unfused glial space, but do not expand further to create the membrane loops observed in *ttx-1* mutants (n=6; Figures 1D, 1F, and 3E), suggesting that *remo-1* is required for both AMsh fusion and AWC expansion. The cell autonomous functions of REMO-1, together with lack of REMO-1 expression in AWC neurons, suggests that the remodeling defects of AWC are non-autonomous, and originate from AMsh glia.

To determine whether *remo-1* regulates other neuronal remodeling events, we examined IL2 sensory neurons, which exhibit extensive dauer-dependent dendritic arborization (Schroeder et al., 2013). We found that 100% of *daf-7(e1372)* animals and 98% of *daf-7(e1372)*; *remo-1(ns250)* double mutants exhibit remodeling, scored using a transgene consisting of an IL2 specific promoter (P*F28A12*.*3*) driving membrane-targeted myristoyl-GFP (myr-GFP) (Figures S2C-S2E). Thus, *remo-1* specifically controls dauer-induced AMsh glia and AWC remodeling.

### REMO-1 differentially regulates *ver-1* transcription and amphid remodeling

To understand how REMO-1 functions with respect to other remodeling genes we previously identified, we examined expression of these genes in *remo-1(ns250)* mutants. We found that expression of neither *ttx-1* nor *ztf-16* is affected (n=150 and n=100 respectively). Furthermore, an AMsh glia-expressed AFF-1::GFP fusion protein remains localized to the anterior portion of AMsh glia in *remo-1(ns250)* mutants (n=98). Thus, *remo-1(n250)* affects expression of P*ver-1*::GFP, but not expression or localization of other genes driving remodeling. Curiously, we found that *remo-1(ns843, ns844*, and *ns845)* predicted loss-of-function deletion alleles, as well as *remo-1(ns841* and *ns842)* premature stop-codon insertion alleles we generated, do not perturb *ver-1* expression upon cultivation at 25°C or in dauer larvae, but still block glia remodeling (Figures S3A-S3D). These observations suggest that the effect on *ver-1* expression is allele specific, and demonstrate that *ver-1* expression can be uncoupled from dauer remodeling. This is consistent with our previous findings that mutations in *ver-1* do not fully block remodeling (Procko et al., 2011). Thus, REMO-1 likely acts upstream of additional remodeling pathways.

### REMO-1-dependent remodeling is required for dauer exit

The purpose of amphid remodeling in dauer animals has been a matter of unresolved speculation. Previous studies demonstrated that the odorant receptor repertoire of amphid sensory neurons is altered in dauer animals (Peckol et al., 2001). This observation raises the possibility that dauers might adjust their sensory tuning to more easily detect nearby favorable environments, which would promote dauer exit. We sought, therefore, to examine whether remodeling-defective animals exhibit dauer exit defects. In dauer animals, pharyngeal pumping, used to suck in and trap bacteria, ceases (Cassada and Russell, 1975; Chow et al., 2006); and resumption of pumping is the first observable sign of dauer exit (Albert and Riddle, 1983). To assess dauer recovery, we therefore exposed animals to GFP-expressing bacteria, which promote dauer exit, and monitored GFP accumulation within the pharynx (Figures 4D and 4E). GFP accumulation coincides with the onset of pharyngeal pumping, as correlated by direct observation (Figure S4A) (Chou et al., 2015; Proudfoot et al., 1993), and can be used to quickly examine many animals at a time. We found that 50% of wild-type animals exit the dauer stage at 2.9 ± 0.27 hrs following introduction to bacterial food, while *ttx-1* mutants, in which dauer-dependent AMsh glia remodeling is defective, exit dauer significantly later, with half of animals accumulating pharyngeal GFP at 4.0 ± 0.06 hrs (Figure 4F). Animals in which AWC neurons are ablated (Beverly et al., 2011), but in which AMsh glia fusion still occurs (Figure S4B), are even more severely delayed than *ttx-1* mutants, with 50% recovering at 4.7 ± 0.06 hrs (Figure 4F). This is consistent with the observation that AWC neurons in *ttx-1* mutants can still expand, albeit in a disorganized fashion, within the confined space delimited by the unexpanded AMsh glia (Procko et al., 2011). These results suggest that remodeling of AWC neurons may drive timely dauer exit. However, *ttx-1* mutants also perturb neuronal temperature sensing, and AWC neuron ablations do not distinguish whether it is the mere presence, or the specific remodeling of AWC neurons, which is important. Therefore, we reasoned that *remo-1* mutants, which fail to remodel both AMsh glia and AWC neurons, and which do not exhibit other obvious defects, might be better suited for testing the effects of remodeling on dauer exit.

We found that following food exposure, 50% of *remo-1(ns250)* animals exit dauer at 4.6 ± 0.56 hrs, a similar delay to that observed in AWC-ablated animals (Figure 4F), supporting the idea that dauer remodeling is indeed required for efficient dauer exit. Importantly, restoring wild-type REMO-1 specifically to AMsh glia of *remo-1(ns250)* animals, not only restores AMsh glia fusion, and presumably AWC remodeling, but fully rescues the dauer exit delay (Figure 4F). Expression of REMO-1 in amphid neurons, however, does not (Figure 4F). Taken together, these results strongly support the notion that dauer remodeling of AMsh glia and AWC facilitates dauer exit.

## DISCUSSION

The studies we present here provide molecular insight into how sensory neurons and their associated glia remodel in response to a changing environment, and demonstrate direct behavioral consequences of such plasticity. Correlating the activities of single neurons, single synapses, or single sensory endings, with animal behavior is essentially impossible in most model systems, as these responses require the activities of large neuronal ensembles. We show here that *C. elegans* provides a unique setting that allows direct interrogation of this black box regime, where single neuron, neuron-glia pair, or sensory structure perturbations can lead to direct behavioral output.

Our data are consistent with a model in which REMO-1 acts in AMsh glia following detection of dauer entry cues to facilitate AMsh glia remodeling. This drives expansion of AWC neuron NREs, which, in turn, promotes efficient dauer exit following re-introduction of *C. elegans* to a favorable environment. REMO-1 functions in parallel to or downstream of DAF-7/TGFβ signaling, and may therefore directly sense environmental cues promoting dauer entry. However, it may also respond to detection of such cues by sensory neurons, become active in response to internal ligands, or exhibit constitutive activity, transducing signals only when downstream signaling machinery becomes available. Future identification of a REMO-1 ligand may help to distinguish among these possibilities. Furthermore, our results suggest that REMO-1 likely has yet to be discovered targets, which may function in both AMsh glia and AWC neurons.

REMO-1 is a member of a small family of predicted *C. elegans* GPCRs with a unique C-terminal sequence. This raises the possibility that these other GPCRs might also participate in remodeling events. Interrogating the consequences of their loss may reveal such roles.

Finally, how REMO-1-mediated expansion of AWC neurons leads to accelerated dauer exit is a fascinating area for future investigation. This study, as well as our previous studies, reveals that AMsh glia fusion occurs in only 50% of wild-type dauer animals. Why this remodeling is not fully penetrant remains unclear. However, it is possible that it represents a way for animals to hedge their bets about the quality of a dauer exit signal. If dauers that remodel exit more quickly, and the environmental stimulus triggering exit is only transient, these animals would not go on to reproduce. However, un-remodeled dauer animals would not initiate dauer exit, and would survive until a bona fide favorable environment is detected. In such a model, the increased surface area of AWC neurons could enhance their sensitivity to favorable environmental stimuli, perhaps by increasing the number of available receptors. Enhanced sensitivity could be used to detect volatiles from further away, or to compensate for the thicker cuticle of dauer animals, which may reduce odorant engagement with receptors. Our studies show that exposure to a dauer-exit stimulus for four hours is sufficient to induce exit of both remodeled and unremodeled animals. Thus, if remodeling is a mechanism for testing stimulus stability, persistence of the stimulus for four hours, but not less than three hours, must be a good predictive measure of longer term stimulus stability in natural contexts.

## ACKNOWLEDGEMENTS

We thank members of the Shaham lab for their insight and comments on the manuscript. Electron microscopy was performed at the Simons Electron Microscopy Center and National Resource for Automated Molecular Microscopy located at the New York Structural Biology Center, supported by grant SF349247 from the Simons Foundation, NYSTAR, and NIH National Institute of General Medical Sciences grant GM103310. Some nematode strains used in this work were provided by the *Caenorhabditis* Genetics Center, which is funded by the NIH National Center for Research Resources, or by the Knockout Consortium. This work was supported by NIH grant R35NS105094 to S.S. and a Women & Science Fellowship to I.H.L.

## AUTHOR CONTRIBUTIONS

I.H.L. and S.S. designed the experiments and wrote the manuscript. I.H.L. performed all experiments and analyses, except for electron microscopy, which was performed by Y.L., and the genetic screen for *ver-1* expression, which was performed by C.P.

