## Supplemental Figures and Methods for "*C. elegans* REMO-1, a glial GPCR, regulates stress-induced nervous system remodeling and behavior"

#### Figure S1

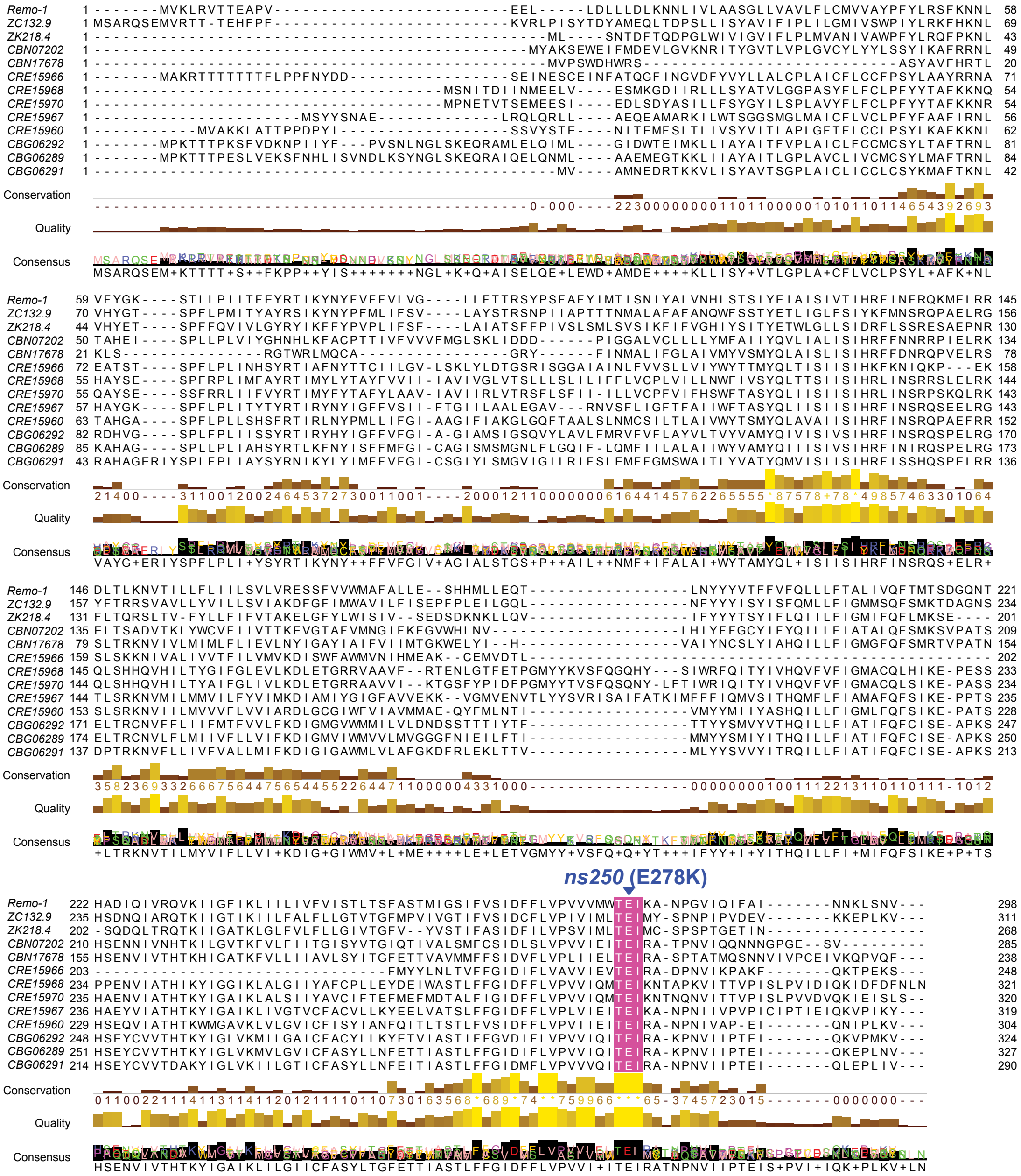


**Figure S1, related to Figure 2. Multiple sequence alignment of the nematode Srz predicted GPCR subfamily.** Sequence alignment using Jalview of *remo-1* paralogs and orthologs in the Srz predicted GPCR subfamily. Paralogs: *ZC132.9* and *ZK218.4*. Orthologs: *CBN07202*, *CBN17678* (*C. brenneri*); *CRE15966*, *CRE15968*, *CRE15970*, *CRE15967*, *CRE15960* (*C. remanei*); *CBG06292*, *CBG06289*, *CBG06291* (*C. briggsae*). *remo-1(ns250)* mutation site is marked by blue arrow. The conserved threonine-glutamic acid-isoleucine (TEI) motif is colored in pink.

FIgure S2

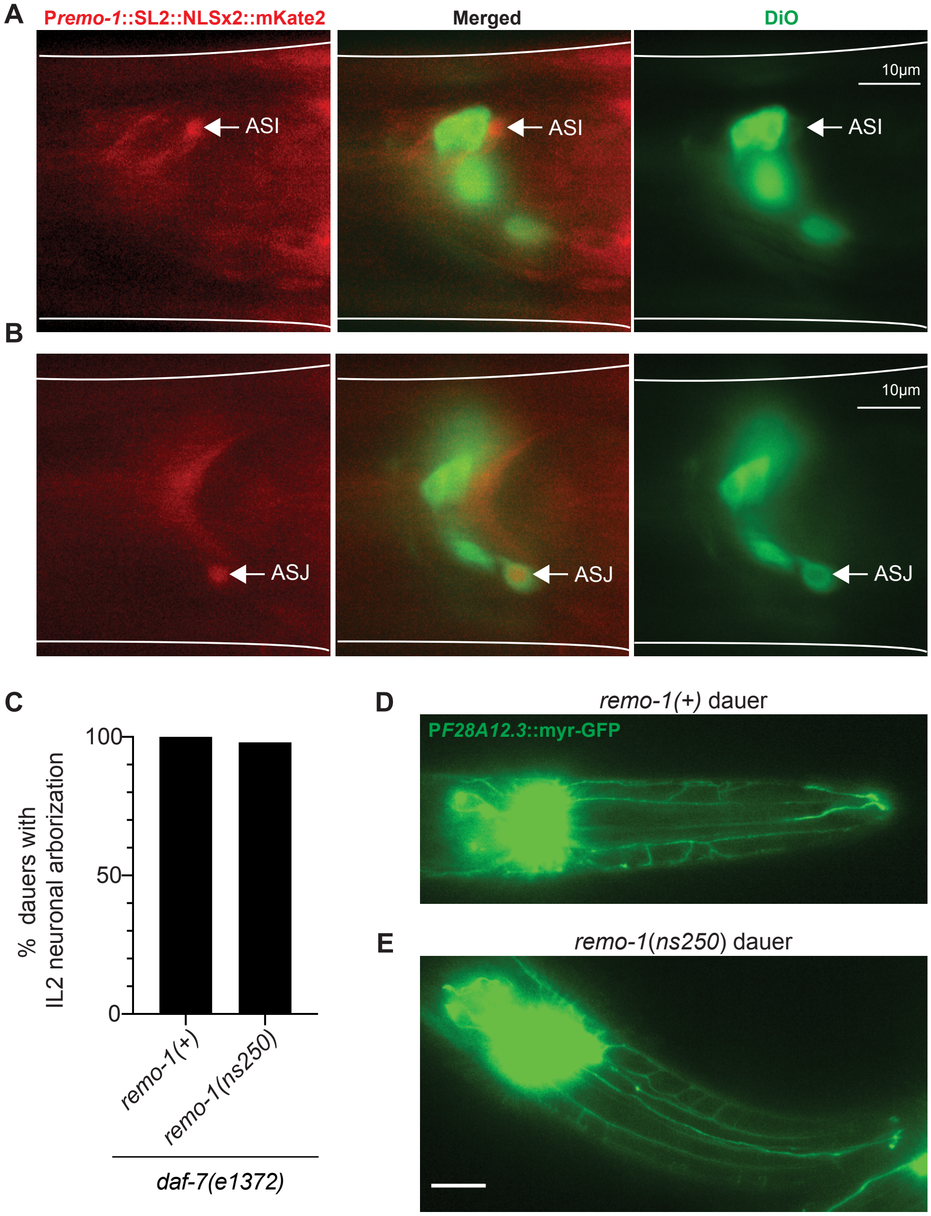


**Figure S2, related to Figure 4. Characterization of *remo-1* expression pattern and effects on other remodeling events in dauer.** (A,B) Fluorescence images of young adult animals carrying *remo-1* transcriptional reporter stained with DiO (right). DiO dye filling assay was conducted to stain amphid sensory neurons *in vivo*. White outlines trace the worm. (A) ASI expression (left); merged (middle). (B) ASJ expression (left); merged (middle). Scale bar, 10µm. (C) Percentage of dauers with IL2 dendritic arborization in indicated genotypes. (D,E) Fluorescence images depicting expression of IL2 reporter (P*F28A12.3*::myr-GFP) in *remo-1(+)* (D) and *remo-1(ns250)* (E) dauer animals. Dauer larvae were induced by the *daf-7(e1372)* mutation cultivated at 25°C. Scale bar, 10µm.

FIgure S3

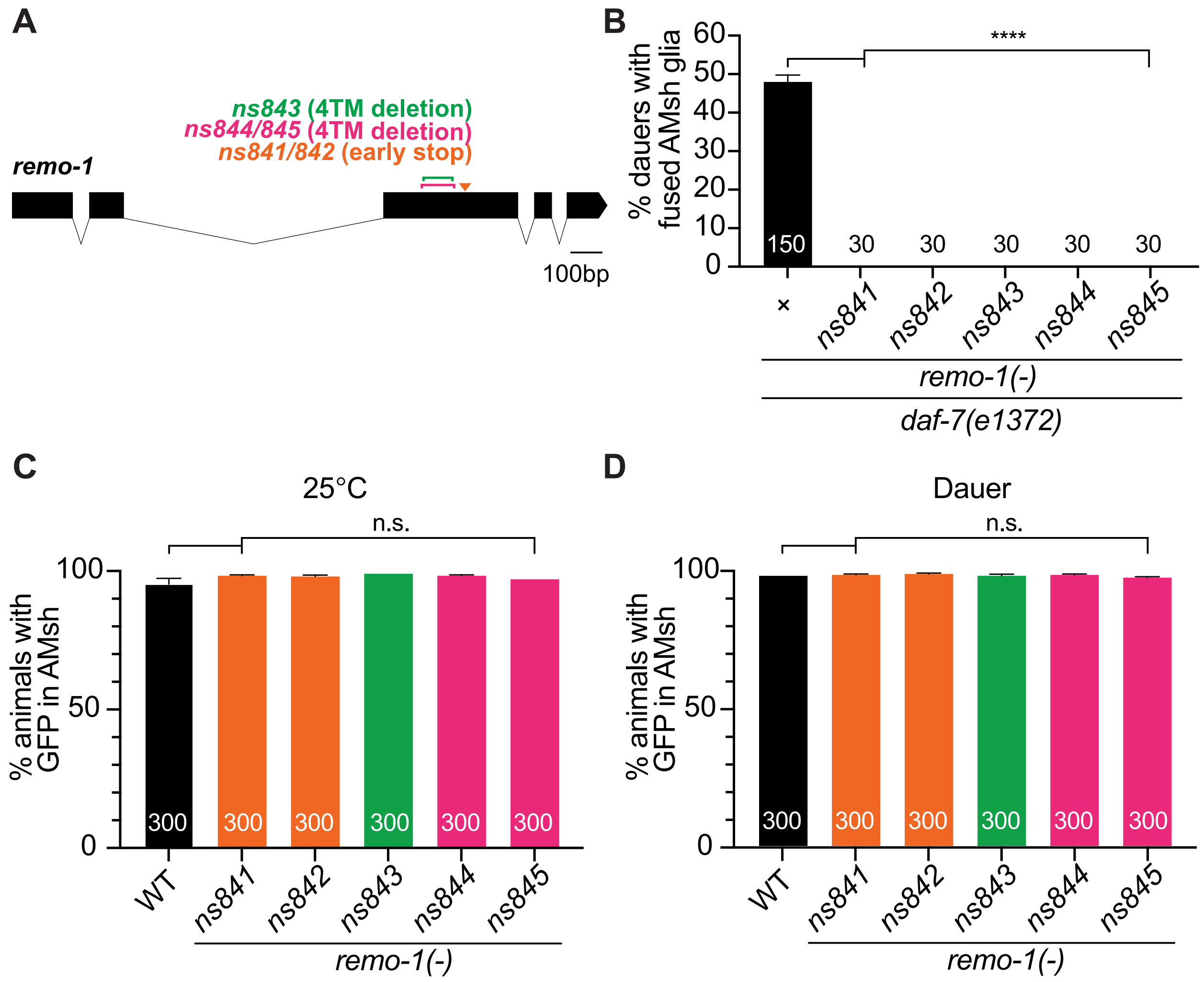


**Figure S3, related to Figure 4. *remo-1* allele genetics.** (A) *remo-1* gene structure and mutation sites. Boxes represent exons and lines indicate introns. Point mutations and deletions relative to the *remo-1* start codon: *ns841/842*: leucine-leucine at positions 185-186 change to alanine-stop codon indicated in orange; *ns843*: 15 amino-acid deletions at positions 157-171 indicated in green; *ns844/845*: 21 amino-acid deletions at positions 154-174 indicated in pink. (B) Histogram detail as in Figure 3B. (C) Histogram detail as in Figure 2A. (D) Histogram detail as in Figure 2B.

Figure S4

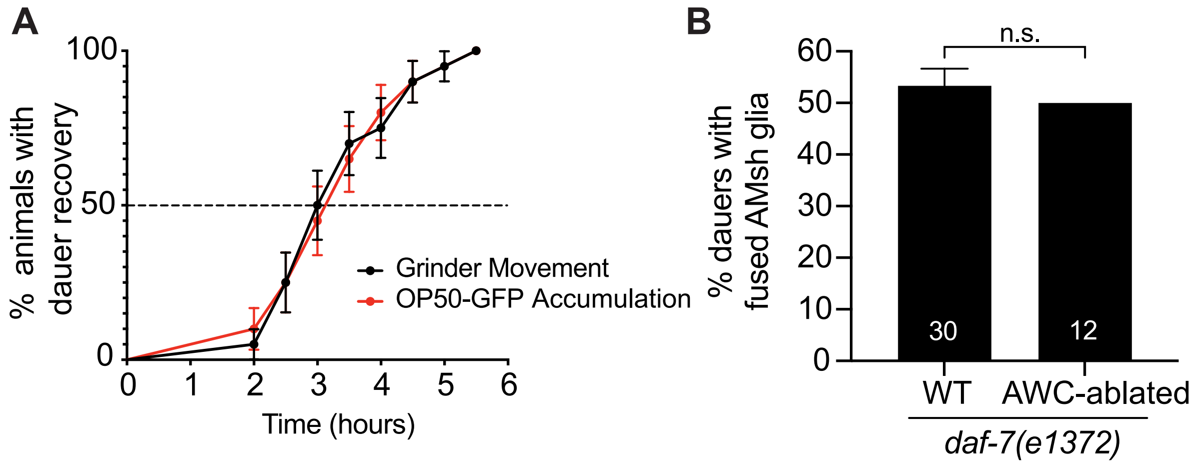


**Figure S4. Dauer exit measured by onset of OP50-GFP accumulation and effects of AWC ablation on remodeling.** (A) OP50-GFP accumulation in the pharynx coincides with the onset of pharyngeal pumping measured by direct observation of consistent grinder movement. Histogram detail as in Figure 4F. (B) Histogram detail as in Figure 3B. n.s., $p > 0.05$.

### MATERIALS AND METHODS

#### LEAD CONTACT

#### MATERIALS AVAILABILITY

Materials generated in this study, including strains, plasmids and clones, are available from the Lead Contact upon request.

#### DATA AND CODE AVAILABILITY

This study did not generate any unique datasets or code.

#### EXPERIMENTAL MODEL AND SUBJECT DETAILS

##### *C. elegans*

*C. elegans* were cultivated using standard methods (Brenner, 1974). All animals were cultivated at 15°C or 25°C, unless otherwise noted. Bristol N2 strain was used as wild-type. Mutants recovered by Ethyl methanesulfonate (EMS) mutagenesis were outcrossed at least three times before use. Most strains had the *nsIs22* (P*ver-1*::GFP) IV reporter transgene. Co-injection markers used were P*unc-122*::GFP, P*unc-122*::RFP, or P*unc-122*::DsRed expressed in coelomocytes, or plasmids as otherwise noted. Integrated transgenic strains were generated with UV/trioxalen treatment (Mello et al., 1991) (Sigma, T6137). AWC neurons were ablated via cell-specific expression of caspases under *ceh-36* promoter sequences (Beverly et al., 2011). Some strains listed below were sourced from the CGC, funded by NIH Office of Research Infrastructure Programs (P40 OD010440).

Alleles used in this work are:
LGIII: *daf-7*(*e1372*), *unc-119*(*ed3*)
LGV: *ttx-1(p767)*, *remo-1(ns250, ns820, ns821, ns841, ns842, ns843, ns844, ns845, ns846*[Venus])
LGX: *ztf-16(ns171)*

#### METHOD DETAILS

##### Forward Genetic Screen

Fourth-stage larvae carrying *nsIs22* in the N2 strain background were mutagenized with ethylmethanesulfonate (EMS) for 4 hours. Individual P0s were picked to separate 9cm NGM agar plates with seeded OP50. F2 animals were screened under a fluorescent dissecting scope (Leica) for mutants that failed to express *ver-1*::GFP in the AMsh glia (Procko et al., 2011; Procko et al., 2012).

##### Gene Identification

A combination of Hawaiian Snip-SNP mapping (Wicks et al., 2001), and whole genome sequencing (Zuryn et al., 2010) using galign (Shaham, 2009) to align reads, was used to identify *remo-1*(*ns250*). The *remo-1* mutation was confirmed by fosmid rescue (WRM0638cF08); *remo-1* genomic transgene rescue; and re-introduction of E278K CRISPR alleles.

##### Germline Transformation

Germline transformation was carried out as previously described (Mello and Fire, 1995). Plasmid mixes containing the plasmid of interest, co-injection markers, and pBluescript were injected into one or both gonads of young adult hermaphrodites (Mello and Fire, 1995). pBluescript was used to adjust the DNA concentration of injection mixes as necessary. Injected animals were singled onto NGM plates and allowed to grow for two generations. Transformed animals were picked onto single plates based on co-injection marker expression, and screened for stable inheritance of the extrachromosomal array. Only lines from different $P_0$ hermaphrodites were considered independent. Isolated strains were staged based on morphology under a dissecting microscope and were screened for P*ver-1*::GFP expression, amphid remodeling, IL2 remodeling and/or dauer recovery.

| Transgene | Construct |
|---|---|
| *nsIs22* | P*ver-1*::GFP (Procko et al., 2011; Procko et al., 2012) |
| *nsEx6181*, *-82*, *-83* | *remo-1* fosmid (WRM0638cF08) (20 ng/µl) + P*unc-122*::RFP (20 ng/µl) + pBluescript (60 ng/µl) |
| *nsEx6184*, *-85*, *-86* | *remo-1* gDNA (50 ng/µl) + P*unc-122*::RFP (20 ng/µl) + pBluescript (30 ng/µl) |
| *nsEx6048*, *6173*, *6174* | P*F16F9.3*::*remo-1* cDNA (40 ng/µl) + P*unc-122*::GFP (20 ng/µl) + pBluescript (40 ng/µl) |
| *nsEx6045*, *-46*, *-47* | P*dyf-7*::*remo-1* cDNA (40 ng/µl) + P*unc-122*::GFP (20 ng/µl) + pBluescript (40 ng/µl) |
| *nsIs391* | P*mir-228*::myr-GFP + *lin-15*(+) |
| *nsEx6249* | pIHL28[P*remo-1*::SL2-NLSx2-mKate2] (90 ng/µl) + P*unc-122*::RFP (10 ng/µl) |
| *nsEx1391* | P*F16F9.3*::GFP + pRF4[*rol-6*(*su1006*)] (Procko et al., 2011; Procko et al., 2012) |
| *nsIs855* | (P*F28A12.3*::myr-GFP) |
| *nsEx1942* | P*ttx-1*::GFP + pRF4[*rol-6*(*su1006*)] (Procko et al., 2011) |
| *nsEx3001* | P*R08E3.4*::GFP + pRF4[*rol-6*(*su1006*)] (Procko et al., 2012) |
| *nsEx2727* | P*F16F9.3*::*aff-1* cDNA-GFP + pRF4[*rol-6*(*su1006*)] (Procko et al., 2011) |
| *nsEx6027* | P*unc-122*::RFP (20 ng/µl) + pBluescript (80 ng/µl) |
| *nsEx6125, -26, -27* | pIL20[P*F16F9.3*::REMO-1 E278K] (10 ng/µl) + P*unc-122*::GFP (20 ng/µl) + pBluescript (70 ng/µl) |
| *oyIs85* | P*ceh-36*::TU#813 + P*ceh-36*::TU#814 + P*srtx-1*::GFP + P*unc-122*::DsRed]. TU#813 and TU#814 are split caspase vectors (Chalfie Lab) subcloned downstream of the *ceh-36* promoter. CGC; (Beverly et al., 2011) |

**Plasmid Construction**

Plasmids were constructed using Gibson cloning into the applicable backbone plasmid (Gibson et al., 2009). *remo-1* cDNA was constructed by cloning each exon from the fosmid WRM0638cF08 and fusing in sequential order using Gibson cloning. *remo-1* promoter region includes the entire *remo-1* genomic locus, including its 5' intergenic region. CRISPR-related vectors were generated using a site-directed mutagenesis protocol on plasmid pDD162 (Dickinson et al., 2013; Liu and Naismith, 2008).

**Additional Plasmids**

| Plasmids | Details |
|---|---|
| pIHL164 | *remo-1*[E278K] CRISPR targeting vector #1, pDD162 backbone |
| pIHL212 | *remo-1*[deletion/premature stop] CRISPR targeting vector #1, pDD162 backbone |
| pIHL213 | *remo-1*[deletion/premature stop] CRISPR targeting vector #2, pDD162 backbone |
| pIHL217 | *remo-1*[Venus] CRISPR targeting vector, pDD162 backbone |

**Cytoplasmic Mixing Assay**

AMsh glia fusion was observed using a fluorescence redistribution assay described in detail in (Procko et al., 2011; Procko et al., 2012). First-stage mosaic larvae carrying the *nsEx1391* array in one of the two AMsh glia were picked onto fresh seeded plates. Mosaic animals were cultivated for at least 48 hours at 25°C to induce dauer entry due facilitated by *daf-7(e1372)* temperature-sensitive allele. Animals were only scored if they were dauer larvae by morphology at the end of the assay period. Mosaic animals were scored for redistribution of GFP to both cells, indicating fusion.

**Dauer Selection**

To induce dauer formation by starvation, embryos were collected by hypochlorite treatment and washed extensively in M9. Approximately 200 embryos were placed onto an NGM plate containing a small lawn of OP50 (~10µL of saturated OP50, 24 hrs before use). Embryos were allowed to hatch and develop at 15°C or 25°C into dauer larva. Failure to include OP50 on the plate resulted in animals that arrested as starved L1 stage larvae. Dauer animals were selected by treatment with 1% SDS in M9 solution for 15 min. Alternatively, young animals carrying the *daf-7(e1372)* mutation were induced to form dauers by incubation at 25°C.

**Dauer Recovery Assay**

Pharyngeal Pumping Assay: Dauer animals induced by starvation were individually selected onto a plate with a thick lawn of OP50 and incubated at 15°C. Dauers were examined every 30-60 min for consistent grinder movement, indicative of pharyngeal pumping, under the dissecting scope.

Uptake of GFP-Expressing Bacteria: OP50-GFP bacteria (CGC) were cultivated in LB media. 24 hrs before the assay, approximately 40 µL of bacteria were plated evenly on the NGM plates and dried overnight in the dark at 15°C. Dauer animals induced by starvation were singled out on the seeded plate and cultivated at 15°C in the dark. Every 30-60 min, dauer animals were screened for GFP accumulation in the pharynx under the dissecting scope.

**Dye Filling Assay**

DiO (3,3′-dioctadecyloxacarbocyanine) (Sigma) stock solution was made at 5 mg/ml in N,N-dimethylformamide. Animals were stained with DiO (stock solution diluted 1:10000 in M9 buffer) for 30–60 min, followed by three M9 washes to remove excess dye.

**Generation of *remo-1* Alleles and Reporter Using CRISPR-Cas9 Genome Editing**

Alleles of *remo-1* were generated using the co-CRISPR-based genome editing method as previously described (Arribere et al., 2014). pDD162 was used as a vector backbone and the following sgRNA sequences were added for each individual CRISPR attempt (E278K: 5’ TCCAGTGGTTGTCATGTGGA 3’; deletion and premature stop alleles: #1 (5’ GCGTTTGTTCAAGCAACATG 3’) and #2 (5’ AATTCTGAGTGTTTTGGTGA 3’)) to generate *remo-1* targeting vectors. A *dpy-10* sgRNA-pDD162-based vector was also generated (5’ GCTACCATAGGCACCACGAG 3’). Singlestranded repair oligos were ordered from Sigma (E279K: 5' TATTTGTGAGTATCGATTTCTTCCTTGTT CCAGTGGTTGTCATGTGGACGaAAATCA AAGCGAACCCGGGTGTTATTCAGATTTT TGCAATTAACAACAAA 3'; *dpy-10*(*cn64*): 5' CACTTGAACTTCAATACGGCAAGATGAG AATGACTGGAAACCGTACCGCATGCGG TGCCTATGGTAGCGGAGCTTCACATGG CTTCAGACCAACAGCCTAT 3') (Arribere et al., 2014). N2 animals were injected with the following mix: 50 ng/µl *dpy-10* sgRNA, 50 ng/µl *remo-1*[E278K] targeting vector (or a mix of 25 ng/µl *remo-1*[deletion/premature stop] targeting vector #1 and 25 ng/µl *remo-1*[deletion/premature stop] targeting vector #2), 20 ng/µl *dpy-10*(*cn64*) repair oligo, and 20 ng/µl *remo-1*[E278K] repair oligo (when appropriate) in 1x injection buffer (20 mM potassium phosphate, 3 mM potassium citrate, 2% PEG, pH 7.5). F1 animals with Dpy or Rol phenotypes were picked to individual plates, indicating a CRISPR-based editing event had occurred. F1 animals were allowed to lay eggs, and then genotyped for successful co-conversion of the *remo-1* locus using PCR and restriction enzyme screening or Cel1 digestion of heteroduplex DNA (Ward, 2015). Non-Rol, non-Dpy F2 animals were then singled and homozygosed for the *remo-1* mutation or deletion.

The Venus reporter was generating by using Cas9-triggered homologous recombination (Dickinson et al., 2013). Venus sgRNA sequence (5' ACGGAAATCAAAGCGAACCC 3') and homologous Venus repair template (5' 1.49kb homolog arm of the 3' end of *remo-1* coding sequence)-*loxP*-reversed *unc-119*(+) cassette-*loxP*-Venus transgene-1.5kb *remo-1* downstream homology arm 3') were used to insert the Venus reporter transgene at the 3' end of *remo-1* sequence. DP38(*unc-119*(*ed3*)) animals were injected with the following mix: 10 ng/µl pGH8 (P*rab-3*::mCherry, Addgene #19359), 5 ng/µl pCFJ104 (P*myo-3*::mCherry, Addgene #19328), 2.5 ng/µl pCFJ90 (P*myo-2*::mCherry, Addgene #19327), 50 ng/µl REMO-1::VENUS targeting vector, 10 ng/µl Venus repair oligo in 1x injection buffer (20 mM potassium phosphate, 3 mM potassium citrate, 2% PEG, pH 7.5). Three injected worms per plate were left to starve for 2 weeks at 25°C. Non-red animals were picked individually and allowed to lay eggs, and then genotyped for successful insertion. The *loxP-unc-119*(+)-*loxP* cassette was subsequently removed by injecting the following mix: 50 ng/µl pDD104 (P*eft-3*::*Cre*::*tbb-2* 3'UTR; Addgene #47551), 2.5 ng/µl pCFJ90 (P*myo-2*::mCherry, Addgene #19327), 47.5 ng/µl pBluescript in EB buffer. The animals in which the *unc-119*(+) gene was excised were isolated and confirmed by DNA sequence analysis. The *unc-119*(*ed3*) mutation was then crossed out. Removing the *unc-119*(+) cassette did not change Venus expression pattern.

**Microscopy and Image Processing**

P*ver-1*::GFP (*nsIs22*) expression was assayed using a fluorescence dissecting microscope (Leica). Young adult hermaphrodites or dauer animals were scored. Compound microscope images were taken on an Axioplan II microscope using an AxioCam CCD camera (Zeiss) and analyzed using the Axiovision software (Zeiss). Some images were collected on a DeltaVision Core imaging system (Applied Precision) with a PlanApo 603/1.42 na or UPlanSApo 60x/1.3 oil-immersion objective and a Photometrics CoolSnap HQ camera (Roper Scientific). Images were deconvolved using softWoRx program (GE Healthcare).

**Electron Microscopy**

Dauer animals for electron microscopy were grown at 25°C and selected for AMsh glial fusion or no fusion based on GFP fluorescence (see cytoplasmic mixing assay for more details). Animals were prepped and sectioned using standard methods (Lundquist et al., 2001). Imaging was acquired with a FEI Tecnai G2 Spirit BioTwin transmission electron microscope equipped with a 4K digital camera. The image acquisition, processing, and analysis was performed by using the SerialEM and IMOD software (Schorb et al., 2019).

**QUANTIFICATION AND STATISTICAL ANALYSIS**

Statistical analysis was performed using GraphPad Prism. Sample sizes and statistical parameters are reported in the main text, figures, and figure legends. When comparing a transgenic line to the parental strain, a minimum of three independent lines were scored. Data is judged to be statistically significant when $p < 0.05$ by unpaired t-test or one-way ANOVA followed by Tukey's test where appropriate. The rates of the onset of GFP-tagged bacteria accumulation in the pharynx were plotted as survival from dauer and were compared by log-rank test using GraphPad Prism 8.

**Protein Structure Prediction**

Prediction of protein structure generated using Protter software (Omasits et al., 2014)

**Multiple Sequence Alignment**

A subfamily of Srz predicted GPCR amino acid sequences were aligned using Jalview v.2.11 (Waterhouse et al., 2009)
